# Isolation of sensory hair cell specific exosomes in human perilymph

**DOI:** 10.1101/2021.04.11.439339

**Authors:** Pei Zhuang, Suiching Phung, Athanasia Warnecke, Alexandra Arambula, Madeleine St. Peter, Mei He, Hinrich Staecker

**Author notes:** Co-corresponding author, Department of Otolaryngology Head and Neck Surgery, University of Kansas Health System, Kansas City, KS 66160, Department of Pharmaceutics, College of Pharmacy, University of Florida, Gainesville, FL 32608.

## Abstract

Evaluation of hearing loss patients using clinical audiometry has been unable to give a definitive cellular or molecular diagnosis, hampering the development of treatments of sensorineural hearing loss. However, biopsy of inner ear tissue without losing residual hearing function for pathologic diagnosis is extremely challenging. In a clinical setting, perilymph can be accessed, so alternative methods for molecular characterization of the inner ear may be developed. Recent approaches to improving inner ear diagnostics have been focusing on the evaluation of the proteomic or miRNA profiles of perilymph. Inspired by recent characterization and classification of many neurodegenerative diseases using exosomes which not only are produced in locally in diseased tissue but are transported beyond the blood brain barrier, we demonstrate the isolation of human inner ear specific exosomes using a novel ultrasensitive immunomagnetic nano pom-poms capture-release approach. Using perilymph samples harvested from surgical procedures, we were able to isolate exosomes from sensorineural hearing loss patients in only 2-5 μL of perilymph. By isolating sensory hair cell derived exosomes through their expression level of myosin VII, we for the first time sample material from hair cells in the living human inner ear. This work sets up the first demonstration of immunomagnetic capture-release nano pom-pom isolated exosomes for liquid biopsy diagnosis of sensorineural hearing loss. With the ability to isolate exosomes derived from different cell types for molecular characterization, this method also can be developed for analyzing exosomal biomarkers from more accessible patient tissue fluids such as plasma.

## Introduction

Sensorineural hearing loss (SNHL) is clinically characterized using the pure tone audiogram combined with speech testing to determine hearing threshold and word understanding. Standard audiometry is a subjective test with techniques built-in to identify malingering and to allow reasonable test retest reliability (Lemkens et al., 2002). Changes in the audiogram do not correlate with loss of a single cell type or the molecular pathology causing the hearing loss (Wu, O’Malley, de Gruttola, & Liberman, 2020). At present, additional physiologic tests (i.e. otoacoustic emissions, evoked potentials) can determine the function of outer hair cells, measure the propagation of the sound signal up the auditory pathway, and analyze the depolarization of inner hair cells. These latter tests are in part only accessible to patients with hearing loss up to moderate to moderate-severe levels. To date we still do not have either a means to biopsy the inner ear to determine the source of pathology and map a treatment course or a blood test to diagnose subtypes of sensorineural hearing loss. Such shortfalls also leave us with an inability to understand the active processes of hearing loss and limits the development of targeted therapeutics for inner ear disease and highlight the need for developing inner ear disease related biomarkers.

Both proteins and non-coding RNAs can be found within the cochlea and could be developed as biomarkers for inner ear disease. Thalman et. al., demonstrated that a variety of perilymph proteins could be identified both in humans and in animal models including ones specific to the cochlea(I Thalmann, 2006; Isolde Thalmann, 2001; Isolde Thalmann, Hughes, Tong, Ornitz, & Thalmann, 2006). Using a mouse model Leary et. al. were able to demonstrate the mouse perilymph proteome could be differentiated from the proteome of cerebrospinal fluid(Swan, Peppi, Chen, & Green, 2009). Comparison of the perilymph proteome of humans, guinea pigs and cats demonstrated that 21 proteins were common to all three species and there was a variance in the number of proteins identified(Palmer et al., 2018). This observation demonstrates that there are differences between humans and animal models that will need to be accounted for when developing biomarkers for inner ear disease. Recent evaluation of patients undergoing cochlear implantation identified 878 protein groups, of which 203 were identified only in perilymph(H. A. Schmitt et al., 2017). Based on these sample group differences in diverse protein groups, proteins related to neurotrophic signaling and cytokines could be identified in individual patients (de Vries et al., 2019; H. Schmitt et al., 2018; Warnecke et al., 2019). We evaluated miRNA expression in perilymph from patients undergoing ear surgery and found markers related to diverse diseases such as otosclerosis and Ménière disease(Shew et al., 2018, 2020; Wichova, Shew, & Staecker, 2019). Additionally, the miRNA profile of perilymph could be correlated to the degree of hearing loss presents (Shew, New, Wichova, Koestler, & Staecker, 2019).

MicroRNA biomarkers for neurodegenerative disorders such as Alzheimer’s disease and even sensory organ diseases such as glaucoma and sudden sensorineural hearing loss have been identified in serum(Dismuke, Challa, Navarro, Stamer, & Liu, 2015; Tanaka et al., 2014). The miRNAs identified in body fluids outside of their cells of origin are transported in exosomes (Chang, Wu, Harn, Lin, & Ding, 2018; Punga & Punga, 2017). Exosomes are small membranous vesicles (~30-150 nm) produced by all living cells from their cellular multi-vesicular bodies in and play essential roles in cell to cell signaling. Other extracellular vesicles (EVs) include microvesicles which are produced from budding off the plasma membrane and apoptotic bodies that are fragments of cells undergoing apoptosis (György & Maguire, 2017; Janas, Sapoń, Janas, Stowell, & Janas, 2016; van Niel, D’Angelo, & Raposo, 2018). Within the inner ear of animal models, exosomes have been identified as a means of protecting hair cells from ototoxic insults (Breglio et al., 2020). Neonatal organ of Corti explants have also been shown to release exosomes altered by an ototoxic stimulus(Wong et al., 2018). Thus, we hypothesize that inner ear specific exosomes could be identified in human inner ear perilymph fluids. However, sampling of human inner ear perilymph fluids to evaluate the presence of exosomes is limited by the precious sample volume (<5 μL), which is only available in limited clinical situations. Herein, we use novel immunomagnetic capture-release nano pom-poms for isolating specific population of hair cell derived exosomes in ultra-low volume of human perilymph, which makes the ultrasensitive detection of exosomal biomarkers possible for precision molecular diagnosis. The developed immunomagnetic nano pom-poms allow the exosome capture via specific antibody from ultra-low sample volume. Under magnetic field aggregation, the captured exosomes can be enriched, and on-demand released through the conjugated photo-click chemistry on the nano pom-poms surface, in turn, harvesting intact exosomes for enhancing the exosomal marker detection specificity and sensitivity(N. He et al., 2021; Zhao, McGill, Gamero-Kubota, & He, 2019). The isolated exosomes exhibit specific expression of protein myosin VII in relation to the status of hearing loss. We foresee this method will enable the ultrasensitive analysis of exosomal biomarkers, potentially from more accessible patient tissue fluids such as plasma for developing non-invasive liquid biopsy diagnosis of sensorineural hearing loss.

## Materials and Methods

### Patients Demographics and Audiometric Evaluation

Adult patients undergoing surgery in which the inner ear was opened were recruited for the study (Table 1). Hearing thresholds were determined for both ears at 0.25, 0.5, 1, 2, 3, 4, 6, and 8 kHz using the descending method of limits (air and bone conduction). Threshold was defined as the lowest audible level, measured twice for that patient. The diagnosis of Meniere disease (Patients 1 and 3) was based on current AAO-HNS guidelines(Goebel, 2016). A CT scan of the temporal bone was performed for all patients which was used to confirm the diagnosis of otosclerosis in patients with conductive/mixed hearing loss (Patients 2 and 5). The diagnosis of progressive genetic sensorineural hearing loss (Patient 4) was based on a family history of multiple female relatives with hearing loss and cochlear implants. For all patients the four-frequency pure tone average (500, 1000, 2000 and 4000 Hz) was determined (based on bone conduction thresholds for the otosclerosis patients).

**Table 1:** Patient demographics

| **PATIENT** | **1** | **2** | **3** | **4** | **5** |
| --- | --- | --- | --- | --- | --- |
| **AGE** | 49 | 50 | 59 | 60 | 41 |
| **GENDER** | Female | Female | Male | Female | Female |
| **DIAGNOSIS** | Meniere Disease | Cochlear Otosclerosis | Meniere Disease | Progressive SNHL (genetic) | Cochlear Otosclerosis |
| **PTA DB)** | 68.75 | 40 | 90 | 50 | 33.75 |

#### Perilymph Collection

Perilymph sampling was approved by the Human Studies Committee. Perilymph collection techniques during surgical procedure that were previously described by our lab were utilized to collect the specimens(Shew et al., 2018). Patients received standard of care treatment other than withdrawal of a 2-5 μl sample using a sterile glass capillary tube (Alpha Laboratories; Eastleigh, Hampshire, United Kingdom). After sample extraction, the procedure was completed in a standard fashion. The samples were placed in a sterile, RNase free tube and frozen at −80° C.

#### Exosome isolation

Perilymph fluids from the inner ear of 5 patients undergoing either cochlear implantation for sensorineural hearing loss (patients 1,3,5) or stapedectomy (patients 2 and 5) for conductive hearing loss and healthy human plasma from one patient provided by KUMC biorepository were analyzed. Exosomes were isolated either using immunomagnetic nano pom-poms via capture-release process, or an adapted sucrose cushion ultracentrifugation method as the control group. The immunomagnetic isolation of nano pom-poms is detailed in our previous publication(M. He, Crow, Roth, Zeng, & Godwin, 2014; Zhang, He, & Zeng, 2016). Briefly, the human perilymph sample was incubated by adding 100 μL nano pom-poms with anti-CD9 conjugation at 4°C overnight. After washing with PBS three times, photo release was performed using an UV transilluminator at 365 nm wavelength at 4 °C for 15 min (~6 mW/cm^2^) to harvest intact isolated exosomes into 100 μL PBS solution. Exosomes from healthy plasma were isolated using sucrose cushion ultracentrifugation. Plasma (1mL) was diluted with 1× phosphate-buffered saline (PBS) to achieve a final volume of 6 mL. The sample was cleared of cell debris and microvesicles by sequential centrifugation at 3000 g for 30 min, and 10,000 g for 30 min at 4 °C. Then, 30% (w/w) sucrose (Fisher Scientific) solution was prepared in 1× PBS and filtered using 0.2 μm syringe filter. The 3 mL sucrose was added to the bottom of each ultracentrifugation tube, followed by gently loading the sample fluid over the sucrose to form two discrete layers. The fluid with sucrose cushion was then spun at 100,000g, at 4 °C for 90 min. Thereafter, the 3 mL sucrose solution was collected from the bottom of the tube and transferred to a new tube. Then 4 mL 1 × PBS was added, followed by gently pipetting the fluid to break up the sucrose bed. Finally, the solution was ultracentrifuged at 100,000g, at 4 °C for 90 min to discard the supernatant and pellet the exosomes.

#### Immune fluorescence nanoparticle tracking analysis

The isolated exosomes from perilymph and plasma were evaluated by nanoparticle tracking analysis (NTA) using a NanoSight NS300 (NanoSight, Malvern Panalytical Ltd, Malvern, UK) and analyzed under both bright field (total particle number) and fluorescence mode (labelled particles with positive surface marker expression). In this study, the anti-Myosin VIIa/MYO7A antibody (Abcam, USA) was used to probe the exosome surface. The isolated exosome samples were diluted (1:4) in 1× cold PBS to a final concentration approximately 5×10^8^ to 1×10^9^ particles/mL (pre-quantified using NTA), followed by incubation with 0.01 μg biotinylated Myosin VIIa antibody at 4°C overnight on a rotator with low rotation speed. Subsequently, 1 μL Streptavidin-Qdot 525 (1 μM, Thermo Fisher) was added to solution and incubated at 4°C for 1h on a rotator. Thereafter, 30 μL of biotin magnetic beads were prepared for each sample (5mg/mL, AbraMag, Thomas Scientific) and washed 3 times using 1× PBS under magnetic field aggregation to remove the supernatant. The washed biotin magnetic beads were resuspended in the exosome solution and incubated at 4°C for 1h on a rotator. After incubation, using a magnetic bar to aggregate the biotin magnetic beads and then collect the EVs for nanoparticle tracking analysis (NTA) under fluorescence mode. The ratio of concentration of exosomes at fluorescence mode to the concentration of exosomes at bright field was calculated as the Myosin VIIa expression level from isolated exosomes in the human perilymph fluid.

#### SEM and immune-staining TEM imaging analysis

Exosome bound beads were resuspended in 200 μL 1× cold PBS solution and washed 2 times with pure water followed by the fixation in a 2% EMS-quality paraformaldehyde aqueous solution for electron microscopic imaging. A 5 μL sample was added to cleaned silicon chips and immobilized after air dry in a ventilation hood. Samples on silicon chips were mounted on a SEM stage by carbon paste. A ~ 5 nm coating of gold-palladium alloy was applied to improve SEM image background. The SEM imaging was performed under low beam energies (7 kV) on a Hitachi SU8230 filed emission scanning electron microscope. For immune staining TEM imaging, a solution of 1×PBS and 0.1% BSA (1mL) was prepared, then a 50 μL of the solution was mixed with ~2.5 μL gold conjugated CD63 antibody (6nM) and the prepared exosome sample grid in a clean 1.5 mL Eppendorf tube. This was incubated for 1 hr. at room temperature. Afterwards, the sample grid was be soaked in 100 μL MilliQ pure water for 30 sec. and transferred onto filter paper to remove all liquid and dried at room temperature for 20 min. Five minutes of negative staining with 2% aqueous uranyl acetate was followed by distilled water washes. Images were acquired at 75 kV using a Hitachi H-8100 transmission electron microscope.

## Results

Using immunomagnetic nano pom-poms for capturing and releasing intact exosomes, we were able to isolate exosomes from ultra-low sample volumes in less than 5 μL of human perilymph, demonstrating the application of magnetic aggregation to collect nano-pom bound exosomes (Figure 1B). The flower-like nano pom-pom structure enables the spatial restriction to bind only the vesicles in size smaller than a few hundred nanometers with affinity capture antibody (here we used anti-CD9 antibody). The exosome bound nano-poms can be aggregated and manipulated via a magnetic field subsequently undergoing washing and light release to separate from nanoparticles on demand. Thus, the harvested intact exosomes can be preconcentrated into a small volume (~100 μL) for downstream molecular analysis.

**Figure1.**
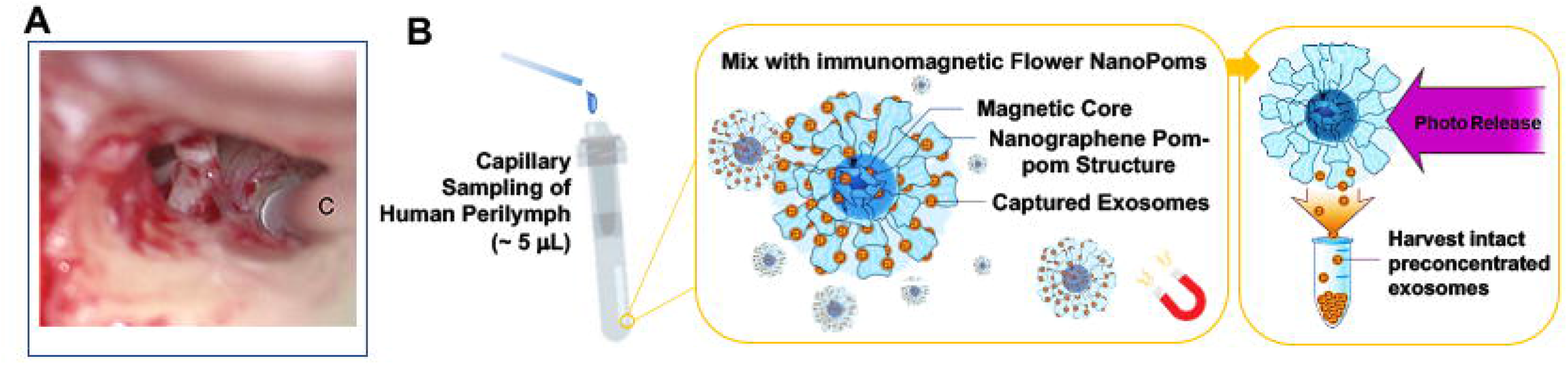
(**A**) Sampling of human perilymph fluids during a right cochlear implant. The capillary (C) can be seen at the round window (**B**) The schematic illustration of immunomagnetic nano pom-poms for capture-release isolation of exosomes from human perilymph in ~5 μL.

The unique flower-like nano pom-poms morphology is shown in Figure 2A which shows the magnetic beads covered with dense nanographene sheet layers for enhanced surface area allowing exosome capture. The perilymph derived exosomes are shown in Figure 2B. The exosomes have a uniform size range below 100 nm which were captured on the surface of immunomagnetic nano pom-poms. We also used immunostaining TEM imaging to confirm the captured vesicular structures as exosomes via probing with immuno-gold (CD63+) nanoparticles (~15 nm). As we can see in Figure 2C, the dense gold nanoparticles in black dots were seen indicating the high amount of capture of perilymph exosomes.

**Figure 2.**
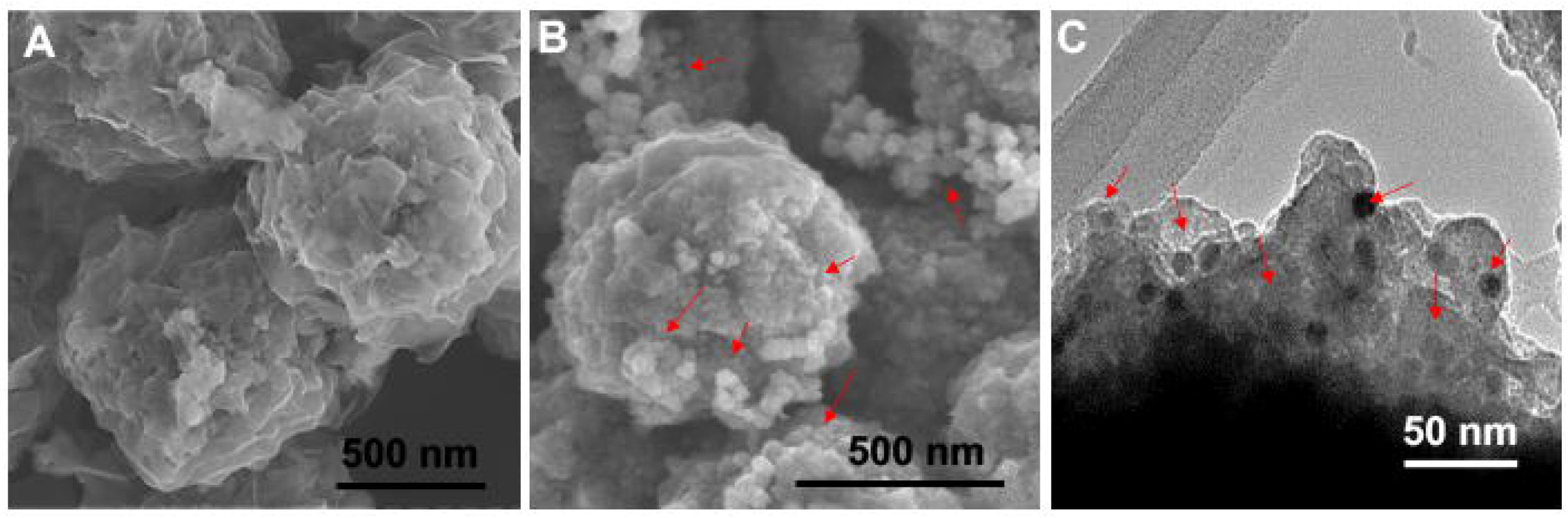
The morphological analysis of immunomagnetic nano pom-pom captured exosomes from human hearing loss patient perilymph fluids. (**A**) The surface property of immunomagnetic nano pom-poms showing the unique flower-like pom-pom nanographene sheet layers conjugated with photo-cleavable antibodies for enhancing exosome capturing performance. (**B**) The perilymph derived exosomes were captured on the immunomagnetic nano pom-poms in the size range around or below 100 nm. The red arrows indicate the abundant clusters of captured exosomes of a uniform size. (**C**) TEM image showing the cross-sectional view of captured exosomes from human perilymph on the surface of immunomagnetic nano pom-poms with conjugated with CD9 capture antibody. The small black dots are the immuno-gold (CD63+) nanoparticles (~15 nm) for probing to confirm exosomes presented on the surface of nano-poms. The red arrows indicate the EV-immuno-gold nanoparticles complexes.

We further used the nanoparticle tracking analysis (NTA) to measure the concentration of isolated perilymph exosomes from three sensorineural hearing loss and two mixed hearing loss patients (Fig 3). Because the sampling process is highly invasive, obtaining healthy control perilymph is impossible. Thus, we used a healthy individual plasma sample to isolate exosomes to serve as the control group. Owing to the specific marker defined exosome isolation using our developed immunomagnetic nano pom-poms, we can extract marker specific (CD9 positive) exosome populations from plasma sample to match perilymph exosome populations, which avoids the interferences from other microvesicles or non-disease associated vesicles. We observed that perilymph contains substantial populations of exosomes in concentrations of ~10^8^ particles/mL, however this is less than what is seen in plasma (~2×10^9^ particles/mL). (Figure 3A). Patients 4 and 5 showed significantly higher concentration of perilymph exosomes than other three patients 1, 2 and 3. To the best of our knowledge, isolating exosomes from human perilymph has not been reported yet. The rat inner ear explant cultured exosomes were reported in the size range from 30-150 nm in a concentration of ~10^8^-10^9^ particles/mL, which is comparable with our observation(Wong et al., 2018). Interestingly, although the dominant size range of isolated exosomes is from 50-150 nm, we observed unique size distribution profiles from each patient perilymph exosomes, which indicates that not all patients have identical size distribution peaks (Figure 3B). Patient 4 and 5 tended to have much more smaller exosomes around 50 nm (~45%) than that (~1%) from patient 1 and healthy control, and the dominant exosome size (~40%) for patient 1 and healthy control is 100-150 nm. It is worth mentioning that we observed exosomes isolated from human plasma exhibiting a completely different profile with higher abundance in larger size (Figure 3B).

**Figure 3.**
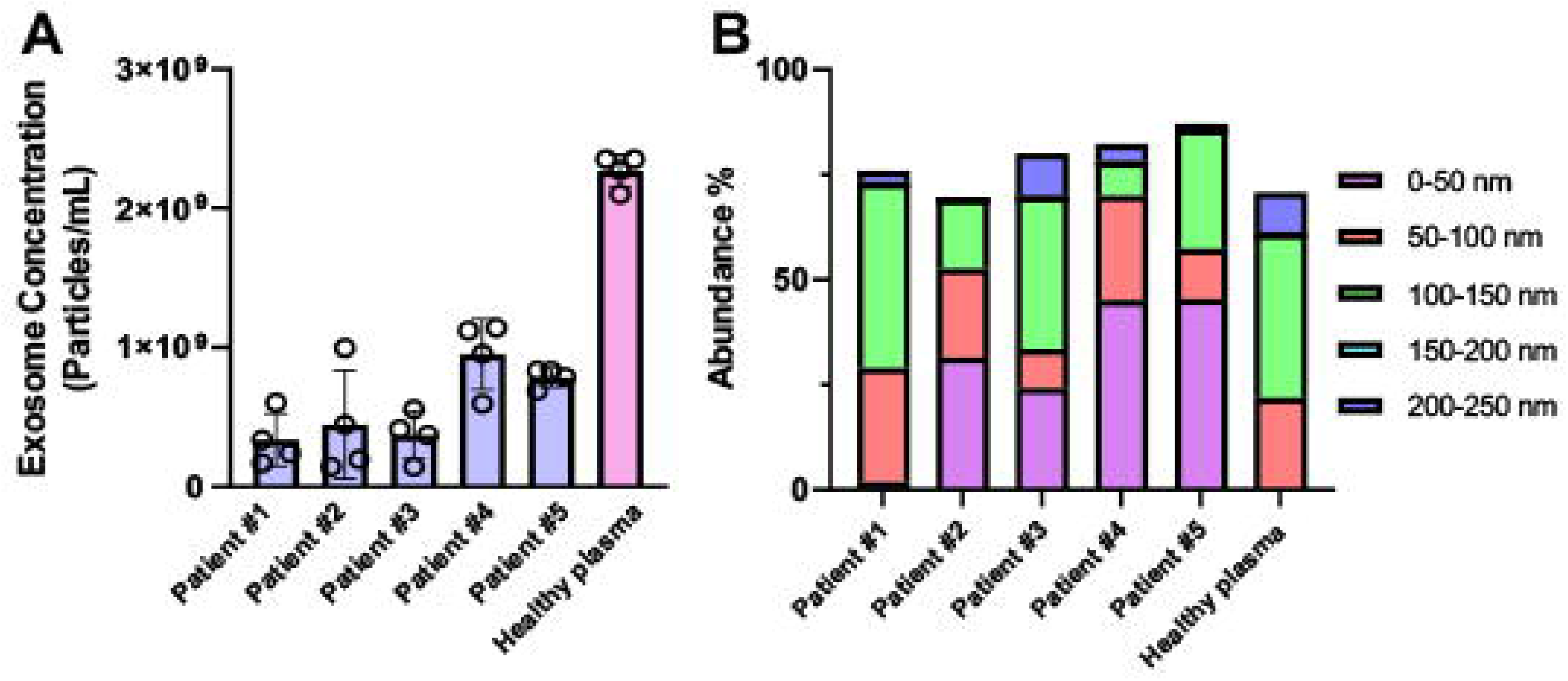
(**A**) The nanoparticle tracking analysis (NTA) for measuring the concentration of NanoPoms isolated exosomes from ~5μL human perilymph collected during surgery. (**B**) The size distribution profiles of isolated exosomes from patient perilymph fluids compared with healthy individual plasma sample using nanoparticle tracking analysis.

Immunolabelling of isolated exosomes for fluorescent NTA analysis enables the characterization of exosomal surface marker expression levels in relation to the total exosomes. The protein myosin VIIa is an essential sensory hair cell specific marker which is highly expressed on hair cells, particularly the actin-rich hair bundle in all species studied(Coffin, Dabdoub, Kelley, & Popper, 2007). Via immunofluorescent NTA analysis, we also observed expression of myosin VIIa on isolated perilymph exosomes (Fig 4). Since detection of fluorescently labelled EVs by NTA is more sensitive than detection of EVs by brightfield, the relative quantification of myosin VIIa positive EVs in relationship to the brightfield NTA EV quantification results in percentages greater than 100% expression in some patients (Fig 4). Using this normalization process we can however compare individual patients. Interestingly, patients 4 and 5 showed much less expression level of myosin VIIa than that patient 1, although the total secreted exosome concentrations from patient 4 and 5 are much higher than that from patient 1. We did not observe any expression of myosin VIIa in healthy plasma isolated exosomes, which is not surprising, as the plasma circulating exosomes may not derive from hair cells. There was no relationship between identifiable patient factors and the percentage of exosomes expressing myosin VII immunoreactivity (Table 1). Even patients with profound hearing loss and decreased caloric responses (Patient 3) demonstrated the presence of myosin VII in perilymph EVs suggesting that hair cells are still present. Interestingly for both Meniere disease (Patient 1 and 3) and cochlear otosclerosis (Patients 2 and 5) the levels of detectable myosin VII were variable when comparing the patients. The patient with the best hearing threshold (Patient 5) actually had the lowest relative expression of myosin VIIa, which could represent the baseline production of EVs by hair cells. Theoretically, perilymph exosomes could be derived either from auditory or vestibular hair cells since all hair cells in the inner ear express myosin VII. The variability in the percentage of EVs expressing myosin VIIa compared to their hearing level (pure tone average) again emphasizes that our understanding of hearing loss is incomplete. Evaluation with additional markers such as prestin may refine our analysis(Parham & Dyhrfjeld-Johnsen, 2016). The present data does demonstrate that we can isolate hair cell specific exosomes from human perilymph.

**Figure 4.**
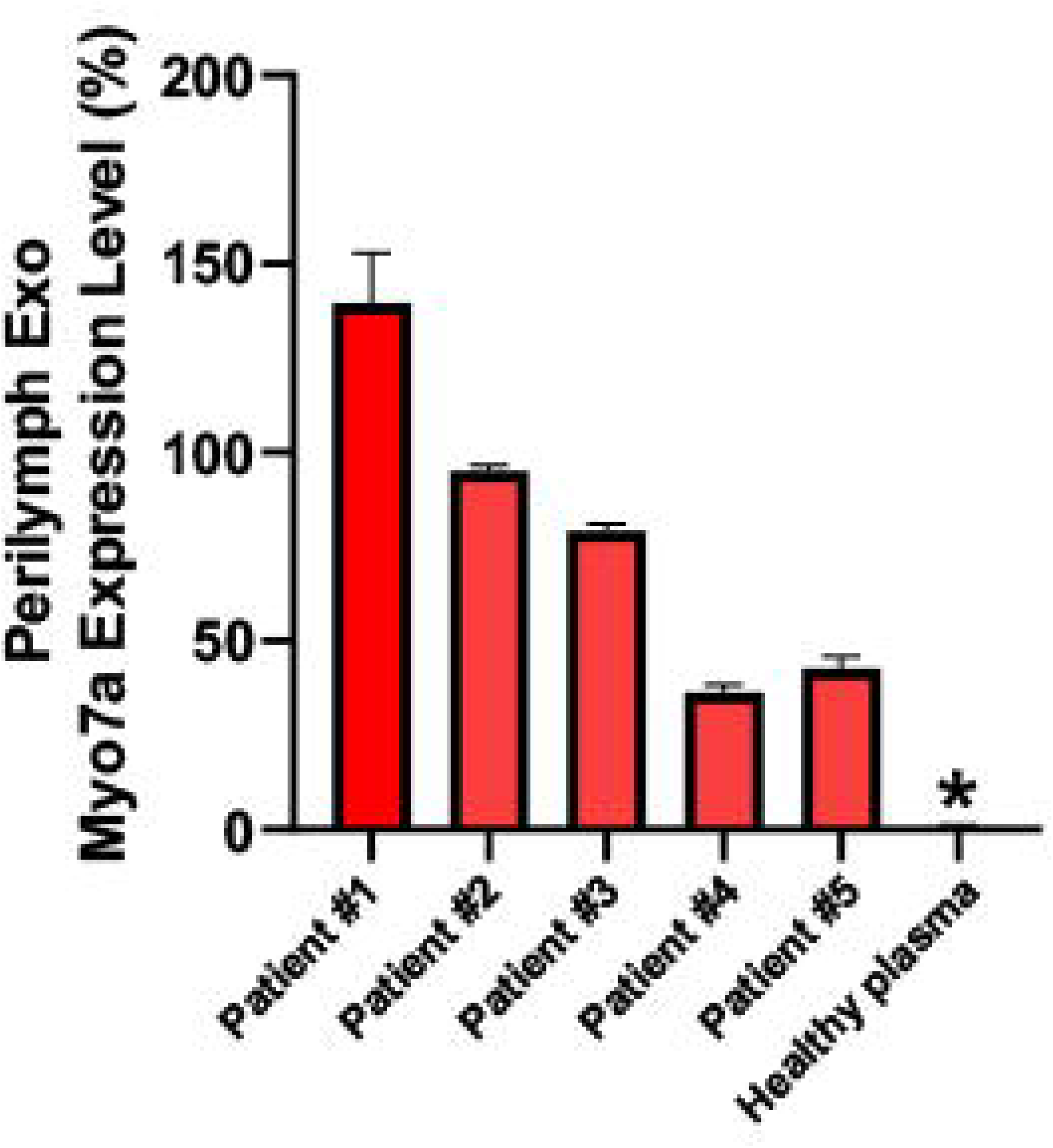
The immunofluorescent NTA analysis of myosin VIIa protein expression levels from isolated exosomes derived from human perilymph fluid collected from patients compared with healthy human plasma exosomes.

## Discussion

Exosomes have been described in all body fluids and are recognized as playing a vital role in intercellular communication in disease and health(van Niel et al., 2018). Using immunomagnetic capture-release nano pom-poms, we for the first time demonstrate the presence of exosomes in human perilymph (Figure 2). The presence of the sensory hair cell specific protein myosin VII on the exosomes suggests that at least a subset of exosomes are produced by either auditory or vestibular hair cells(Naples, Miller, Ramsey, delivery, & translational, 2019). Diagnosis of inner ear disease on a molecular or cellular level presents unique challenges due our inability to date to biopsy the inner ear and the complex tissues and multiple pathophysiologic processes involved in hearing loss. Since exosomes are transported from their tissue of origin and have been evaluated as disease biomarkers for a range of different disorders. Like the central nervous system (CNS), the inner ear is separated from the circulation by a blood-labyrinthine barrier that inner ear derived exosomes would have to cross(Banks et al., 2020; Bodmer et al., 2002; Matsumoto, Stewart, Banks, & Zhang, 2018). Numerous studies in neurodegenerative and neurological disorders have identified plasma exosomes that can be associated with prognostic or diagnostic outcomes. (De Toro, Herschlik, Waldner, & Mongini, 2015; Leggio et al., 2017). Progress is also being made in the identification of serum exosomes related to eye disease which as an organ system shares many of the properties of the inner ear (complex neuroepithelium, difficult to access and biopsy)(Klingeborn, Dismuke, Bowes Rickman, & Stamer, 2017). A common strategy for matching CNS based exosomes to disease process when taking a serum sample is to analyze the exosomes protein or miRNA payload which has been shown to be altered by disease states(Huang et al., 2018). At present we do not have a complete characterization of the proteome or miRNAs associated with inner ear disease states due to current limitation in accessing the inner ear for purely diagnostic purposes. Additionally, the small size of the inner ear (160 μl fluid volume) and limited number of sensory cell population (30,000 auditory sensory hair cells) make it likely that a more sensitive method is required to capture inner ear related exosomes from serum.

The developed immunomagnetic nano pom-poms are uniquely position for isolating exosomes from precious volume of human perilymph fluids, owning to the highly specific exosome capture-release process and magnetic field aggregation for exosome enrichment which is particularly essential for handling ultra-low sample volume. In this work, we demonstrated first time to be able to isolate perilymph exosomes and probe the specific myosin VIIa hair cell markers from isolated perilymph exosomes, which is much more sensitive than ultracentrifugation exosome isolation approach(N. He et al., 2021).

Inner ear specific proteins have been found in the serum of animal models of ototoxicity and in human vestibular disease(Liba et al., 2017; Mulry & Parham, 2019; Parham et al., 2019). Serum miRNA associated with sudden sensorineural hearing loss have also been found in humans (Li et al., 2017; Nunez, Wijesinghe, Nabi, Yeh, & Garnis, 2020). It is presently not known either of these is associated with exosomes that have been transported out of the inner ear. A diverse population of inner ear cell specific that could potentially be used to select exosomes specific to the inner ear from the serum population followed by either proteomic or PCR analysis of their cargo. The ability to capture exosomes from the serum based on inner ear specific markers could also open the possibility to diagnose inner ear disease on a cellular level since many of the inner ear specific proteins are expressed by particular cell types (Naples et al., 2019). Further studies will be needed in patients with acute and chronic hearing loss to further translate these findings.

### Conclusion

Using immunomagnetic capture-release nano pom-poms, the current study provides proof of concept that human inner ear exosomes could be identified in microliter ultra-low sample volume from patients undergoing surgery where the inner ear is opened. Additionally, we could identify exosomes that were derived from sensory hair cells. As in CNS based neurodegenerative diseases, exosomes potentially could provide a novel path towards developing inner ear diagnostics.

## Abbreviations

(CNS): central nervous system
(EV): extracellular vesicles
(miRNA): micro RNA
(SNHL): sensorineural hearing loss

## Notes

### Competing Interest Statement

The authors have declared no competing interest.

